# Photoelectrochemical imaging of action potentials for single cardiomyocytes through contact force manipulation of organoids

**DOI:** 10.1101/2022.09.05.506608

**Authors:** Rachel Jacques, Bo Zhou, Emilie Marhuenda, Jon Gorecki, Anirban Das, Thomas Iskratsch, Steffi Krause

## Abstract

Accurate monitoring of cardiomyocyte action potentials (APs) is essential to understand disease propagation and for trials of novel therapeutics. Patch clamp techniques offer ‘gold standard’ measurements in this field, but are notoriously difficult to operate and only provide measurements of a single cell. Here we propose photoelectrochemical imaging (PEI), in conjunction with a setup for controlling the contact force between the cardiomyocyte organoids and the sensor surface, for measuring APs with high sensitivity. The method was validated through the use of drugs, and the results successfully visualized the expected electrophysiological changes to the APs. PEI allows for several cells to be monitored simultaneously, opening further research to the electrophysiological interactions of adjoining cells. This method expands the applications of PEI to three-dimensional geometries and provides the fields of stem cell research, drug trials and heart disease modelling with an invaluable tool to further investigate the role of APs.

## Main

Sudden cardiac death (SCD) kills more than 100,000 people per year in the UK alone [1]. The precursor for SCD is arrythmia arising from improper ion channel properties and function, which can be caused through a number of diseases both inherited and acquired [2]. It is for this reason that accurate measurements of cardiomyocyte (CM) action potentials (APs), *in vitro*, is extremely important, both to follow disease propagation and to effectively trial potential therapies. However, current measurement techniques suffer from low throughput or lack of ability to measure in higher ordered tissue models, thus limiting their usability.

Current ‘gold standard’ measurements of cardiomyocyte APs are acquired through manual patch clamp assays. This technique, although providing high throughput and direct measurement of the APs, is extremely complex to operate and can only measure one cell at a time [2], [3]. Numerous optical techniques have been implemented through the use of dyes that are voltage sensitive or through genetic editing to express voltage sensitive proteins [4], [5]. Although these methods allow for multiple cell measurements simultaneously, they only provide relative voltage changes and can only be used for short periods of time due to the toxic nature of the markers and photobleaching. Multi-electrode arrays (MEA) have also been used to measure the extracellular field potential (FP) of cardiomyocyte monolayers cultured in multi-well plates [6], [7]. The FP is a direct result of the AP but differs in shape and so the use of MEA leads to loss in information of the underlying ion channel mechanisms. More recently a MEA has been coupled with electroporation to achieve greater coupling efficiency, thereby allowing for direct AP measurements to be obtained [8]. Simultaneous measurements can be taken at different points across a syncytium within a single well, but again individual cell measurements cannot be obtained using this method. The technique is limited as individual AP morphologies and temporal changes cannot be distinguished on a cellular basis. The method is also limited in its ability to measure 3D geometries, which are of particular interest in both drug screening and disease propagation modelling due to increased cell maturity [9], [10].

Light-addressable potentiometric sensors (LAPS) are a photoelectrochemical imaging technique, which allows for photocurrent measurements at electrolyte-insulator-semiconductor (EIS) structures. A bias voltage is applied to the structure to create a depletion layer at the semiconductor-insulator interface, and a scanning modulated light source, which is focused within the semiconductor layer, induces an alternating photocurrent. Changes in charge on the insulator surface give rise to local changes in the width of the depletion layer, and therefore changes in the measured photocurrent [11], [12]. LAPS measurements have been used extensively to monitor local ion concentrations, impedance and pH [13], [14].

Traditionally LAPS have been unable to measure the charge of the cell membrane due to the charge screening effect. This effect arises due to the small gap between the sensor surface and cells in cultured media, which is filled with electrolyte and is larger than the Debye length of the electrolyte (typically < 1nm in physiological media), i.e. charges from the cell membrane are ‘screened’ from the sensor [12], [15]. This is demonstrated in many experiments where a very low signal strength was present from the membrane charges [14], [16]. For example, in an experiment where the membrane potential of aplasia cells was measured using LAPS, although the measurements correlated with that of the membrane potential, the change measured represented only 1.3% of that in the membrane [15].

LAPS has previously been used to measure cardiac APs with a silicon substrate [17]. The cardiomyocyte cells were cultured on the sensor surface and light was focused from the front side, i.e. through the cells. Although it was claimed to successfully measure and monitor drug induced changes in the APs, the method has since been debunked and the changes measured are thought to be due to mechanical movements of the cells [18]. The mechanical movement and shape changes are a direct result of the AP and would result in alterations in the focus of the light source and so changes in the photocurrent over time.

To maximise the speed of photoelectrochemical measurements, silicon remains an attractive choice of semiconductor due to its narrow bandgap and high charge carrier mobility, which is compatible with high modulation frequencies required for high-speed imaging. This is particularly useful when attempting measurements of fast cellular responses, such as APs. Recently, ultrathin insulators such as self-assembled monolayers (SAMs) have been immobilized on silicon on sapphire (SOS) substrates thereby significantly increasing the photocurrent contrast and sensitivity of LAPS [19], [20]. This material was successful in the imaging of cell surface charges of multiple layers of yeast Saccharomyces cerevisiae [21]. In this case the cells were imaged in agarose gel to reduce the distance between the cells and the insulator surface. However, single cell measurements were not achieved as the cell-surface contact was poorly controlled by the agarose gel.

Here, we present a novel method for LAPS measurements of CM APs from a 3D organoid structure using a SAM-modified SOS substrate. Individual CM organoids (Figure 1a) are pressed to the sensor surface with a well-defined force to achieve excellent contact between the cells and the LAPS device (Figures 1b and c). In this way, the enhanced sensitivity of the substrate is exploited, whilst the problematic focal distance of cells in standard culture can be avoided [22], [23], [24]. Measurements using this technique are compared to standard 2D culture of CMs on the sensor surface. The method is further validated by monitoring the effects of several drugs on AP duration and amplitude. Our study using the force probe-modified LAPS setup, has demonstrated excellent sensitivity to CM APs and their drug responses. The technique has the ability to monitor adjoining cells simultaneously and has opened up the potential of future applications in 3D in vitro models for drug testing, or implementation for *in vivo* measurements.

**Figure 1.**
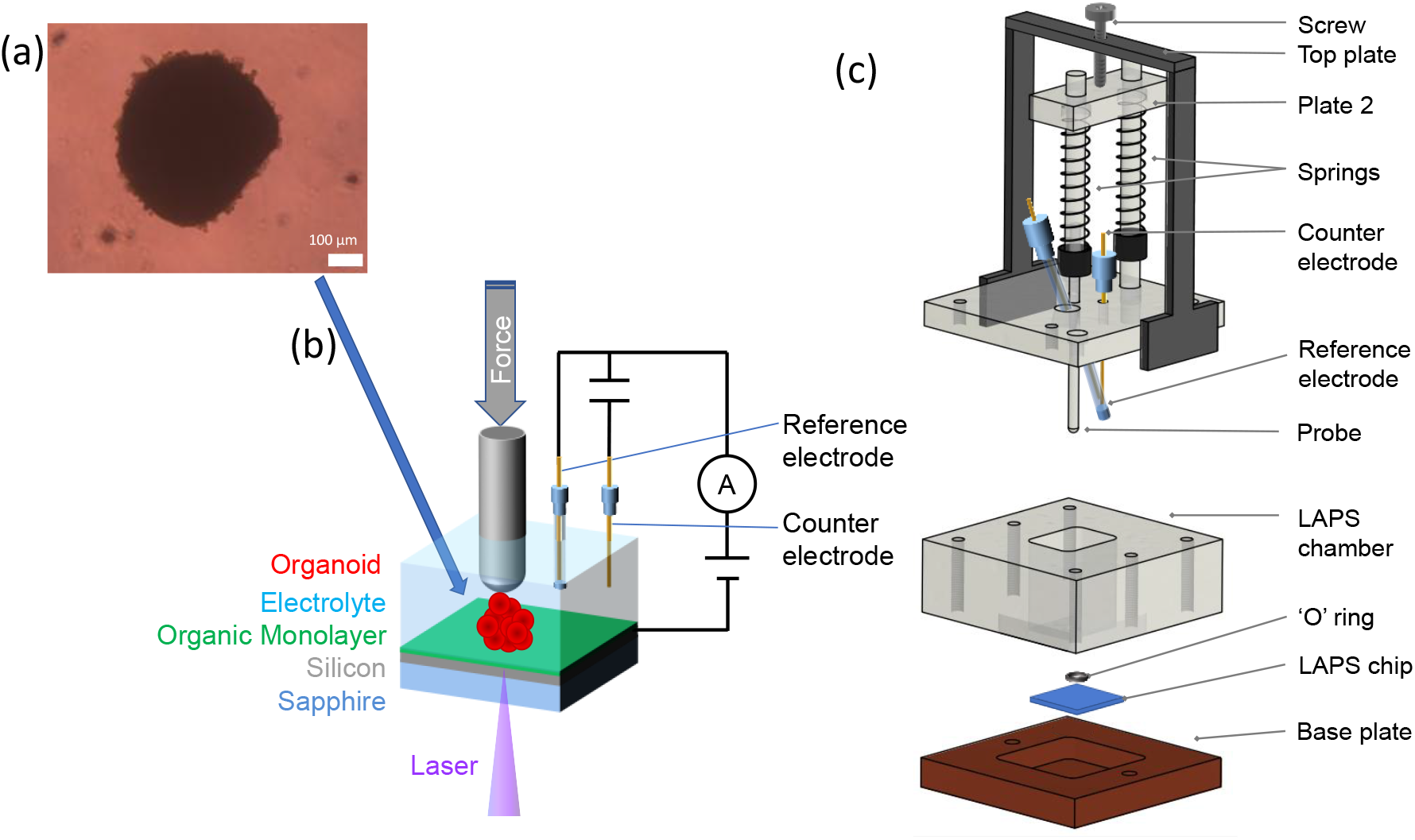
Photoelectrochemical imaging of organoids pressed onto the sensor with a force probe. (a) an organoid image taken using an optical microscope (b) the schematic of the photoelectrochemical system; a scanning modulated 405 nm laser is focused on the silicon layer through the sapphire. An organic SAM is used as an insulating layer between the semiconductor and electrolyte. A CM organoid is positioned on the sensor surface and a force probe is used to apply pressure to the organoid and achieve good contact between cells and sensor surface. (c) the schematic of the force probe assembly together with the LAPS chamber, silicon substrate and base plate. As a screw is positioned and screwed through the top plate, plate 2 is pushed down, compressing the springs and exerting an equal force upon the probe acting downward. In this way, the contact force of the organoid on the sensor surface can be controlled and maintained during measurement.

## Results

### Effect of increasing contact force on cell to the sensor surface

The photoelectrochemical imaging technique, LAPS, has been limited to measurements of 2D cell culture on the sensor surface. Natural cell adhesion focal points ensure a small gap between the sensor and surface [25], [26]; in silicon-based LAPS, this shields the cell membrane charges from the sensor, making imaging and monitoring of the action potential unviable. In addition, culturing cells on a LAPS chip has limited appeal as there is little or no control over the mechanical properties and the surface morphology of the semiconductor substrate. 3D in vitro models have increasingly gained popularity for drug testing and discovery science, due to the enhanced functional maturity and similarity to in vivo tissues, for applications from cancer research to infectious disease modelling. Likewise, investigating mechanically active and sensitive cells such as cardiomyocytes (CMs) in a 3D organoid has the advantage that CMs retain their contacts in all directions, preserving more of the complex chemical and mechanical interplay and communications between cells, thus creating a more realistic model for *in vitro* research [27]. Here, we use a method to ensure good mechanical contact between cell and sensor through careful control of the contact force and the use of 3D organoids (Figure 1).

Initially, the sensitivity of the sensor was established in standard 2D cell culture by photocurrent imaging (Figure 2b) and by monitoring the photocurrent at a fixed position under a cell (Figure 2c). Thinning of the insulating layer using anodic oxides (6.7 nm thick) has been shown to improve sensitivity in these sensors, when compared to thermal oxides (70 nm thick) [28]. In the case of SAMs the layer is even thinner, enhancing sensitivity further [19]. Despite the enhanced sensitivity, no individual cells could be distinguished in the photocurrent image of CMs in culture (Figure 2b). Several cells were monitored with time and the photocurrent amplitude was compared to that of cells with increased contact force through pressing of an organoid onto the sensor surface (Figures 2d and e). Typical clinical heart catheters measure contact forces below 0.4 N [22], and significant damage to cardiac tissues is inflicted at forces above 0.2-0.3 N [23], [24]. We were aiming to apply the minimum force required to achieve good contact with the sensor. Hence, all organoid measurements were taken using a force of 0.02 N.

**Figure 2.**
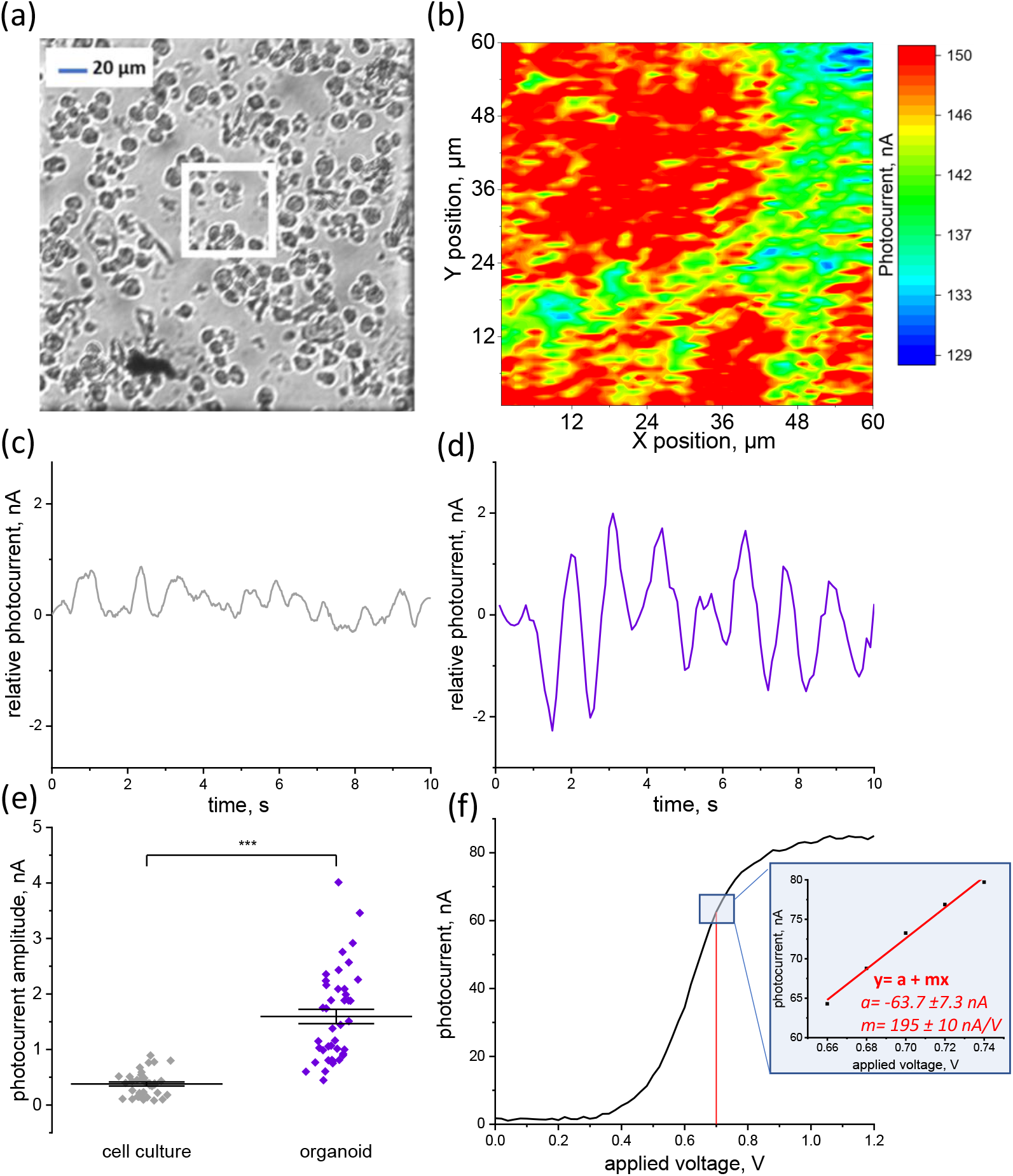
Comparison of photocurrent data of CMs cultured on the sensor surface and organoids pressed onto the sensor with a defined force. (a) an optical image of the cardiomyocyte cells cultured on the LAPS sensor surface highlighting the area of the corresponding LAPS image (white square) (b) the corresponding LAPS image (c) shows the time dependence of the relative photocurrent (photocurrent – average photocurrent) for the cells cultured on the LAPS surface (d) the time dependence of the relative photocurrent for cells of an organoid pressed to the sensor surface (e) superplot of the amplitude of cells cultured on the surface compared with those of an organoid pressed to the sensor surface. The peak to trough amplitude is plotted for each beat of the signal for each cell measured, and the mean and SEM are shown. (f) a typical photocurrent vs voltage curve for the SOS SAM substrate at 20 kHz laser modulation with a zoomed in extract at the 0.7 V location giving a straight line fit for the data along with the parameters used.

The comparison in the acquired signal both in the raw data shown in Figure 2c and in the averaged data (Figure 2d) depicts a clear (over 4.2 times) increase in photocurrent amplitude for cells in the organoids that have a well-defined contact force. Figure 2d also shows a clearer periodic signal indicating the AP of the cell has been recorded when sufficient contact has been achieved. The average amplitudes and approximate frequencies for each cell measured are detailed in Table S3. A typical photocurrent-voltage curve (Figure 2f) was used to establish an approximate translation of the changes in photocurrent (amplitude in nA) to changes in potential (amplitude in mV); this allows for a direct comparison with the signals measured using the ‘gold standard’ patch clamp methods. Using the photocurrent-voltage curve (Figure 2f), an approximate straight line can be fitted to the data at the position of 0.7 V (operating bias in all the measurements). In this case, a change in photocurrent of 1 nA is approximately equal to a change in voltage of 5.1 mV. From this, we can determine that on average the photocurrent measurements with increased contact force have an overall amplitude of approximately 8.1 mV.

### LAPS monitoring of Verapamil effects in the action potential

Verapamil is a calcium ion channel blocker and is used to treat certain cardiac rhythmical defects and to lower pressure in the blood [29]. Here we compare the recorded APs of cells within the organoid before and after adding verapamil to the electrolyte.

LAPS images were collected of cells that have sufficiently good contact to the sensor. In contrast to LAPS images of cells in culture (Figure 2b) individual cells in the organoid could clearly be identified by a local minimum in the photocurrent (Figure 3a). The decrease in the photocurrent is caused by the negative surface charge of the cell membrane [21]. The distance between the minima of the photocurrent is in agreement with the assumption that the cells are adjoining considering the typical sizes of cardiomyocytes in Figure 2a. An appropriate line scan through the cells, shown in Figure 3a as a black line, was continuously recorded to build up a time dependent photocurrent signal for each cell before and after verapamil was added (Figures 3c). The results show a reduction in frequency and amplitude for both cells but also highlight a key change in their phases. Before verapamil is added, the cells are 165 ± 11 ° out of phase with one another as determined by the sinusoidal fit method (see Figure S3 and Table S1). After verapamil is added, the cells’ APs become much more synchronised and are now only 7.37 ±10.7 ° out of phase (see table S4 for a further pair of adjoining cells). The averaged beat profile for each cell (Figure 3b) allows for visualisation of the changes in the AP as a result of verapamil. The period and amplitude of the APs can be determined and compared (Figure 3d). Verapamil significantly increased the duration of each part of the AP (Table S2) and causes a reduction in the amplitude of the APs

**Figure 3.**
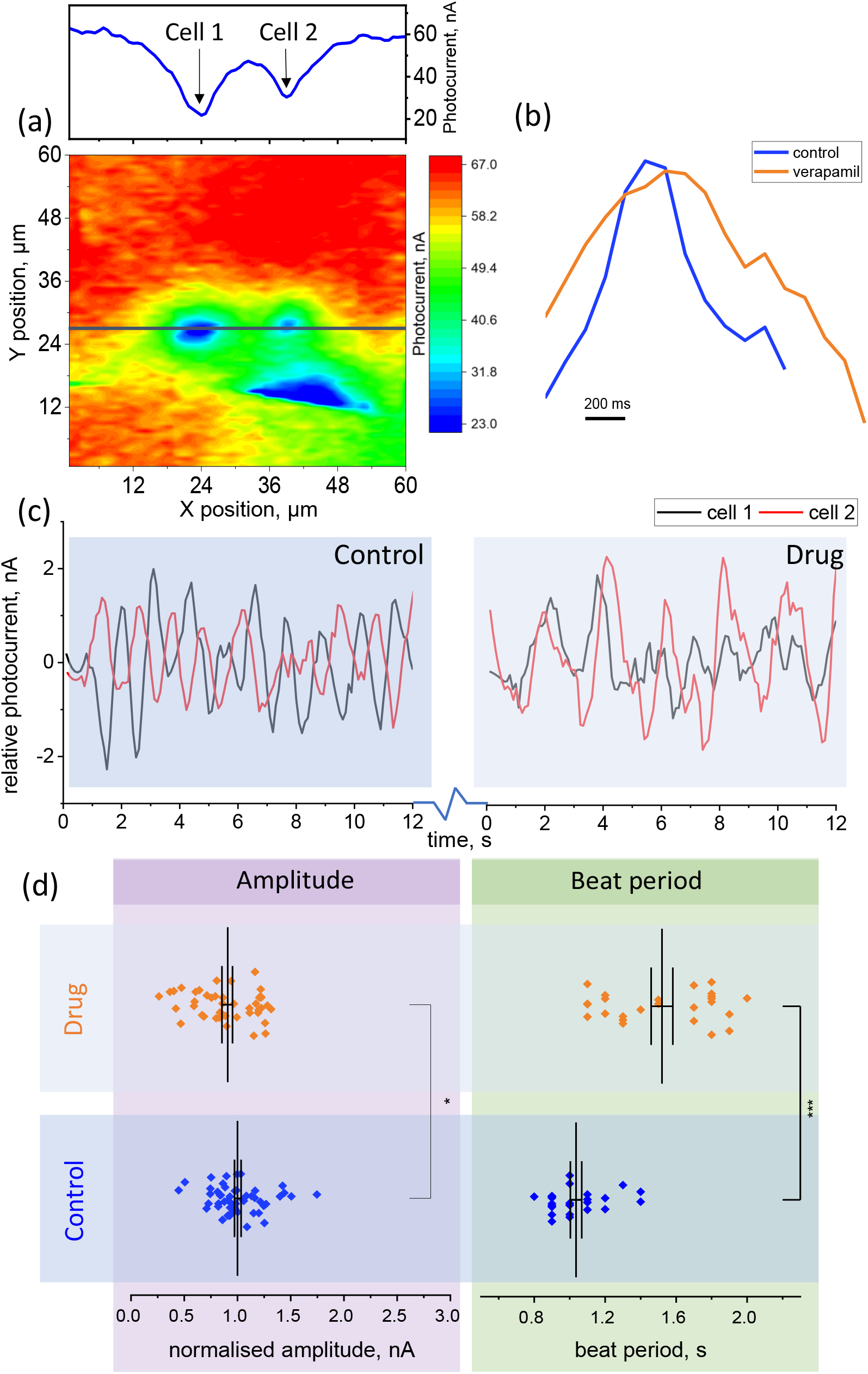
The effect of verapamil on the action potential of individual CMs in an organoid (a) the LAPS image of 2 adjoining cardiomyocyte cells and the corresponding line scan through the data, highlighting the position of the line scans for cell 1 and cell 2; (b) the beat profile before and after verapamil is added averaged over 5 beats. (c) the photocurrent time dependence for cells 1 and 2 before verapamil is added, showing clear phase difference, and 10 mins after verapamil has been added, the cells now show less phase difference. (d) superplots of the amplitude and beat period for before and after verapamil is added showing mean and standard error in mean (SEM) values and the statistical significance of the change.

### LAPS monitoring of phenylephrine (PE) effects on the action potential

PE increases the frequency of the APs and is commonly used to treat hypotension [30], [31]. In Figure 4a a single cell is visible in the LAPS image; the line scan was run continuously at the position of the black line in Figure 4a to build up a photocurrent vs time signal before and after the addition of PE (Figure 4c). The results show an increase in frequency after PE is added; this change is also highlighted in the shortened BP (see averaged AP profile of a single cell Figure 4b) This drug also results in an increase in amplitude for all cells measured (Figure 4d). There was no significant effect of PE on the phase shift between the APs of two adjoining cells (Table S5). The average durations, amplitude changes and phase shifts are detailed in Table S2.

**Figure 4.**
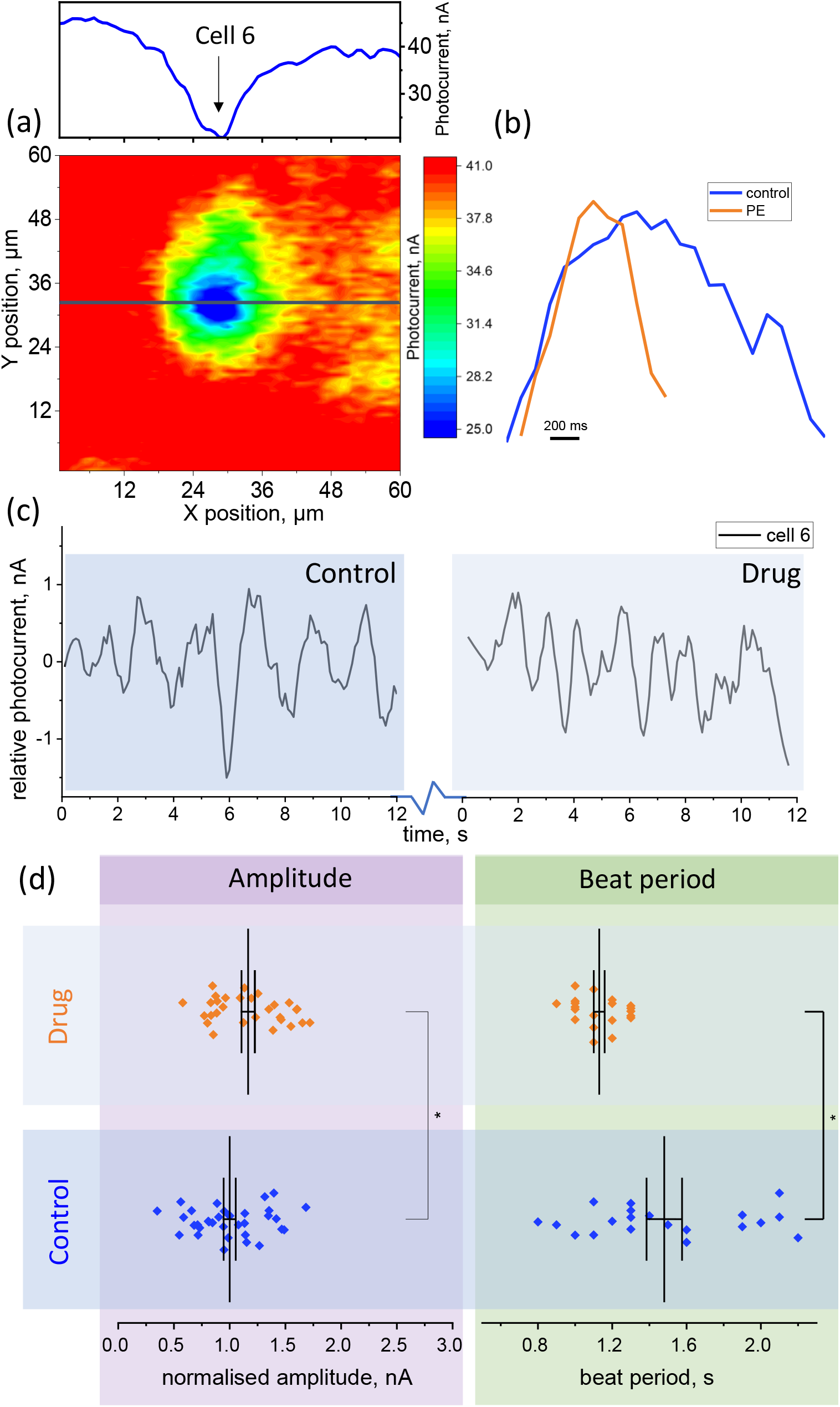
The effect of phenylephrine on the action potential of individual CMs in an organoid (a) the LAPS image of a cardiomyocyte cell and the corresponding line scan through the data, highlighting the position of the line scans for the cell (b) the beat profile of the cell before and after PE is added averaged over 5 beats. (c) the photocurrent time dependence for the cell before and 2-5 mins after PE is added. (d) superplots of the amplitude and beat period (BP) for before and after PE is added showing the mean and SEM values and the statistical significance of the changes

### LAPS monitoring of blebbistatin effects on the action potential

Blebbistatin inhibits the sarcomere from contracting and thus stops the CM cells beating [32]. In this case photocurrent vs time signals were acquired for before and 10 mins after blebbistatin was added (Figure 5). The same analysis as for the other drugs is completed by establishing beat profiles for each cell before and after blebbistatin is added (Figure 5b). In this case, there appears to be a small decrease in all durations and an increase in the amplitude of the APs (Figure 5d), however, none of the changes were determined to be statistically significant.

**Figure 5.**
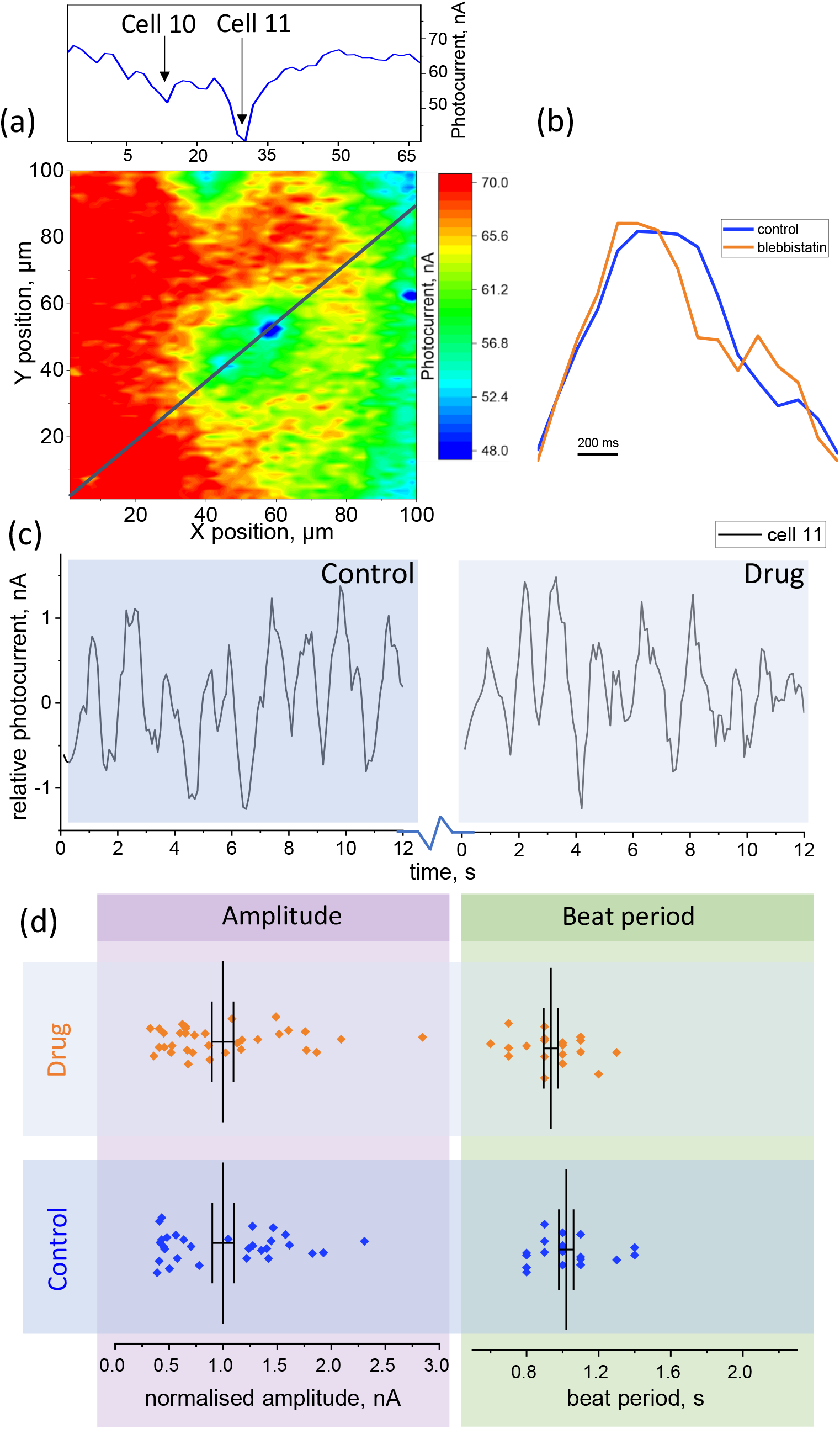
The effect of blebbistatin on the action potential of individual CMs in an organoid (a) the LAPS image of cardiomyocyte cells and the corresponding line scan through the data, highlighting the position of the line scans for the cells (b) the beat profile of ‘ cell 1’ before and after blebbistatin is added averaged over 5 beats. (c) the photocurrent time dependence for the cell before and 10 mins after blebbistatin is added. (. (d) superplots of the amplitude and beat period (BP) before and after blebbistatin is added showing mean and SEM values with no statistically significant changes

## Discussion

The initial results using the new technique, combining LAPS with enhanced cell-sensor contact through defined pressure on organoids compared with LAPS in standard cell culture shows a vastly improved signal strength. The technique displayed its ability to image cells and monitor dynamic potential changes in the cell membrane. The large increase in amplitude of the measurements in Figure 2e highlights these advancements through the improved signal to noise ratio. Standard morphology of neonatal CM (N-CM) APs may however be lost through the sampling interval and through subsequent smoothing of the signal to remove high frequency noise (Figure S2). This is shown through the lack of multi-phasic repolarization culminating in the missing ‘notch’ seen in typical AP measurements [33]. Despite this discrepancy, the clear periodic nature of the signal as well as the changes in the beat profile arising from the drugs used, leaves us in no doubt that the APs of N-CMs have successfully been measured using this technique. Cells with an increased contact force have an average AP amplitude of 8.1 mV, as determined by the line of best fit described in Figure 2f, with some measurements achieving amplitudes as high as 13.8 mV. Measurements of action potentials taken using patch clamp techniques see amplitudes of approximately 80-100 mV [33]–[35]. Increasing the contact force has therefore drastically improved AP signal acquisition using LAPS from the previously reported 1.3% [15] of the AP to about 10-15%

Verapamil has previously been reported to reduce contraction and beat rates [29]. This effect is shown in the reduction of amplitude in Figure 3d for all cells measured. However, a much greater difference is achieved in those cells which display a higher initial amplitude (Table S4). As described previously, the initial amplitude largely relies on the contact between cell and sensor; most likely cells with a smaller amplitude had a lower contact force and so their absolute changes in amplitude also appeared smaller. In Figure 3d the beat period is significantly increased after the addition of verapamil, complying with the expected effect of this drug in slowing of the APs. The APs after verapamil has taken effect show significant lengthening to the AP morphology at all points across the beat profile (Table S2) [36]. In the case of adjoining cells, the phase of the two cells is clearly synchronised when the drug takes effect as can be visualised through comparison in Figure 3c. A second 2-cell comparison for verapamil can be seen in the SI (Table S4) showing the same effect, i.e. bringing the two cells’ APs into a synchronised signal. Clearly, more experiments and results need to be obtained to conclude any additional effects of the drug on a cell-to-cell basis but these initial findings show the potential of the new technique in providing new insights to the field.

PE has been shown to increase both the amplitude of the AP and the rate of beating [30]. In the results shown in Figure 4(d), it is clear that both the amplitude and AP frequency significantly increased as expected. Blebbistatin has no direct effect on the electrophysiology of the CMs but instead inhibits contraction of the sarcomere through inhibition of myosin II ATPase activity, [32], [37]. Its use in this experiment acts as a control to prove that the LAPS measurements are monitoring the APs and not any secondary effects arising from the mechanical movements of the cells. The results show no significant change in amplitude or AP duration in Figures 5b and d. As blebbistatin has restricted any mechanical beating, the photocurrent peaks measured after addition of the drug can only be due to the electrophysiological change in the cells’ membrane arising from their APs

This new method of photoelectrochemical measurements combined with a well-defined force to establish good contact between cells and sensor is able to successfully image and monitor CM APs on a cellular basis. A clear improvement compared to previous cell culture methods can be seen and several drugs have tested the system to determine the reliability of the results obtained. The expected electrophysiological drug induced changes to the AP were successfully visualized with this method. The added advantage of this technique is the ability to monitor several cells simultaneously to within the limit of the scan range (160 μm for the current setup). This will have great effect on the analysis of novel therapies as well as understanding of disease propagation as further knowledge of adjoining cell interactions is critical in this pursuit. Furthermore, the label-free and non-invasive nature of LAPS allows for long-term measuring and ensures that the cells are not disturbed from their natural behaviors. In contrast to patch clamp, LAPS is easy to use and can be integrated with standard microscopes and may also be adapted for in vivo applications in the future. Further experiments of fast cellular responses will now be possible through the advantages of using silicon as this method has proven to overcome the ‘screening’ effect. This novel method has also expanded the applications of photoelectrochemical imaging techniques to 3D cell structures providing new applications to the field.

## Methods

### Materials and chemicals

Silicon on sapphire (SOS) with a 0.5 μm thick silicon (1 0 0) layer (boron doped, 0.05 Ω cm) on 475 μm thick sapphire was used as sensor substrate (Monocrystal, Russia). All other chemicals, unless otherwise stated, were purchased from Sigma-Aldrich and used as received. 1,8-nonadiyne (98%) was redistilled from sodium borohydride (99% +) and stored under argon

### Substrate preparation

SOS samples were prepared as previously reported [20]. In short, SOS was cut to 5 mm x 5 mm pieces. Ohmic contacts were established on the samples by thermal evaporation to coat 30 mm Cr and 150 mm Au on the edge of the chip, before heating to 300 °C for 5 min on a hotplate. The substrate was placed into piranha solution ((3:1) H_2_SO_4_(96%):H_2_O_2_(30%)) at 100 °C for 30 min and then rinsed thoroughly with copious ultrapure water to clean it. The clean substrate was then etched to achieve H termination using 2.5% HF solution for 90 s. The sample was rinsed with ultrapure water, blow dried with nitrogen and placed into a Schlenk tube under vacuum before redistilled 1,8-nonadiyne was added and degassed by several freeze-pump-thaw cycles until no gas bubbles evolved from the solution. The reaction mixture was kept under an argon atmosphere and left for 3 h at 165 °C. Once cooled to room temperature, the substrate was rinsed with DCM and blown dry with nitrogen.

Further functionalization was achieved through a copper catalyzed azide alkyne cycloaddition (CuAAC) or “click” reaction with a slight modification to the reported procedure [20] using a 6-azido-hexanoic acid. The sample was transferred to a “click” solution containing the 6-azido-hexanoic acid (1 ml, 15 mM, ethanol/water 2:1), copper(II) sulfate pentahydrate (1.1 mol % relative to the azide), sodium ascorbate (10 mol % relative to the azide) and TMEDA (0.45 mM). The reaction was completed over 24 h in the dark. The unreacted reagents were removed from the sample by rinsing with ultrapure water and ethanol and blown dry with nitrogen. The treated substrates were then stored in a nitrogen box prior to use.

### Cell and organoid culture

Neonatal rat cardiomyocytes (NRC) isolation and primary cell culture were realized as described previously by Pandey P. et al [38]. Neonatal rat hearts were extracted, carved, kept in ice-cold ADS buffer (116 mM NaCl, 20 mM Hepes, 0.8 mM NaH_2_PO_4_ 5.6 mM glucose, 5.4 mM KCL, 0.8 mM MgSO_4_) and washed once with fresh ADS buffer. Then heart pieces were incubated with 5 ml of enzyme solution (ES, 0.6 mg pancreatin and 246 U collagenase / ml in ADS), for 5 min at 37 °C, under strong shaking and supernatant was discarded in order to wash out red blood cells. This was followed by 5 to 6 digestion runs until the heart pieces were digested. Each time, 5 ml of fresh ES was added to the hearts, incubated 15 min at 37 °C, under vigorous shaking and pipetted up and down 30 times. After each time, the ES containing cells was transferred into plating medium (65% DMEM, 17% M199, 10% horse serum, 5% fetal bovine serum (FBS), 2% glutamax, 1% penicillin/streptamycin (P/S)). Digests were combined in one tube with plating medium, filtered through a 100 μm cell strainer to remove undigested tissue and centrifuged at 1200 rpm for 5 min at room temperature, before resuspended in plating medium. NRCs were pre-plated for 90 min to enrich the cardiomyocytes and remove potential fibroblasts. Cardiomyocytes were then deposited into 96-well plates round bottom coated with polyHEMA to avoid adhesion. The polyHEMA was prepared by adding 5 g of it into 500 mL of ethanol 96%, and 200 μl were deposited into each round bottom well. The plate was then left overnight at 60 °C to leave the ethanol evaporated. Then 10000 cells were incubated for 5 days into 300 μL of medium until formation of a cohesive spheroid in the round bottom wells. Plating medium was used at day one post isolation, and then changed for maintenance medium (maintenance medium 77% DMEM, 18% M199, 2% horse Serum, 2% glutamax, 1% P/S) for four subsequent days until spheroid formation. The beating is shown in Video S1.

For the standard cell culture on the sensor surface, the substrate and LAPS chamber were thoroughly cleaned (with ethanol at 70% for 20min before being rinsed with sterile water and left under UVs for 20 min) and assembled before the surface was coated with fibronectin at 1μg/ml in PBS for 1h at 37°C and washed twice with PBS. Dissociated cardiomyocytes (10000 cells) or spheroids were deposited within the chamber containing plating medium. The media was changed to maintenance media after 24 h and was changed every 2-3 days. Measurements were taken after 4-7 days after seeding once an actively beating monolayer was achieved.

### LAPS set up

The setup for LAPS measurements has been described previously [39]. In short, scanning of the laser is achieved through the use of a MEMS mirror, the amplitude of which sets the scan area. The 405 nm (max 50 mW) laser is modulated at a frequency of 20 kHz and focused onto the silicon from the back of the SAM modified SOS substrate (Figure 1b). As silicon has a strong absorbance at this wavelength, any purely optical effects originating from the interaction of light with the cells can be completely discounted. A bias voltage of 0.7 V is applied between the substrate and an Ag/AgCl electrode to obtain a depletion layer at the silicon-insulator interface. The LAPS chamber was placed in a Faraday cage.

The organoid is positioned onto the sensor surface within the LAPS chamber using a pipette and 1 mL of maintenance medium is added. The force probe is then assembled as described in Figure 1c and the probe is carefully lowered onto the organoid before half a turn is given to the screw through the top panel. This pushes plate 2 downward, compressing the springs by 2.5 mm. The springs having constant k = 0.008 N/mm (Lee Spring LP 008A 06 S316) then apply a contact force of 0.02 N upon the organoid according to Hooke’s law. The probe itself is a 2 mm diameter acrylic rod with a silicone hemisphere of the same diameter on its tip. The hemisphere is set from clear silicone (151 silicone sealant). The minimum force which resulted in accurate and consistent AP measurements was determined to be 0.02 N. Using this force allows for reliable contact with minimal disturbance of the mechanical beating of cells. Beyond 0.1 N some organoids sustained breakdowns in their cell connections with irreversibly damage to the spherical structure of the organoid.

### LAPS measurements

For the 2D cell culture an optical view, using a camera, allowed for accurate positioning of the laser spot on a cell which can be seen to beat. Once a position was decided the photocurrent at the spot was recorded with a sample rate of 0.01 s.

For imaging and recording of cardiomyocyte cells in an organoid several LAPS images were taken at varying positions beneath the organoid, with the aid of optical view. The white light for the optical view was directed through the organoid and so the images were unable to resolve cells in contact with the sensor. The optical view provided the outline of the organoid and so gave an area in which to search with LAPS images.

Once a suitable cell or adjoining cells had been identified by local minima in the photocurrent, a line scan was positioned through the center of the cell or cells. Each line took 0.1 s to complete and, by successive line scan recording, a time profile was established. A detailed description how line scan positions were chosen is provided in the SI (Figure S1).

### Drug dosing of organoids

An initial LAPS scan was taken as described above and then drugs were added to the electrolyte in the following concentrations: verapamil 10 μM, phenylepherine 1 mM, and blebbistatin 50 μM.

Another LAPS scan was then taken after an approximate interval of 2-5mins for PE and 10 mins for verapamil and blebbistatin to allow the drugs to diffuse thoroughly through the electrolyte and take full effect within the cells.

### Data analysis

The time vs photocurrent data for all measurements is smoothed using the Savitzky-Golay method at 50 pts for the cell culture data and 10 pts for the line scans of the cells in the organoid (Figure S3). This rejects the high frequency data resulting from the noise of the LAPS system. All photocurrent vs time plots have their baseline (mean value) subtracted to obtain the relative photocurrent changes over time.

Amplitude for the cell culture and organoid cells is displayed as the average peak to trough decrease in photocurrent during the repolarization phase. The average amplitude before each drug is added is used to normalize all peak to trough values for each cell as the starting amplitude has a large variation depending on the initial contact between cell and sensor. These normalized amplitudes are used in the T-test to determine the significance of the changes. 5 beats from each recording of the photocurrent are used to determine the beat period, (1) rise time (take off point to the peak of the AP), (2) action potential duration (APD), taken from the beat start to 30%, 50% and 90% of the repolarisation (100%= peak to trough) and (3) beat period (BP) taken as the time between the start of each period. The different duration parameters are illustrated in Figure S4. The time dependence data for the organoids is then averaged over 5 beats to create an AP profile of each cell before and after drugs are added. Each beat is normalized so that it ranges from 0 to 1 before they are averaged.

All changes displayed in the paper are statistically significant, unless otherwise stated, using T-test, having p-values of less than 0.05. The significance of changes in amplitude and beat period are indicated on the superplots for each drug with *(p<0.05) **(p<0.01) and ***(p<0.001). The significance of all other parameter changes is listed in the SI (Table S2).

## Supporting information

Video of beating organoid

## Acknowledgements

The authors are grateful to QMUL for providing a PhD studentship to RJ and to EPSRC (EP/R035571/1, EP/V047523/1) for funding.

## Author contributions

RJ: Force probe design, sample preparation, LAPS measurements, data analysis and interpretation, writing-original draft reparation, reviewing and editing. BZ: experimental design, sample preparation, reviewing and editing. EM: cell and organoid culture, reviewing and editing. JG: data analysis and presentation, reviewing and editing. AD: measurement software, selection of measurement parameters, data analysis, reviewing and editing. TI: experimental design, analysis and interpretation of data, reviewing and editing. SK: experimental design, project administration and supervision, funding acquisition, data analysis and interpretation, reviewing and editing.

## References

[1] “What is the most common cause of sudden cardiac death? | Patient.” https://patient.info/news-and-features/what-is-the-most-common-cause-of-sudden-cardiac-death (accessed Mar. 30, 2022).

[2] Hossein. Ardehali, Roberto. Bolli, and Douglas. Losordo, Manual of research techniques in cardiovascular medicine. pp. 50–59 2014.

[3] J. Liu and P. H. Backx, Cardiac Tissue Engineering: Methods and Protocols, Methods Mol Biol, vol. 1181, pp. 203–214, 2014, doi: 10.1007/978-1-4939-1047-2_18.

[4] M. P. Hortigon-Vinagre, V. Zamora, F. L. Burton, J. Green, G. A. Gintant, and G. L. Smith, “The Use of Ratiometric Fluorescence Measurements of the Voltage Sensitive Dye Di-4-ANEPPS to Examine Action Potential Characteristics and Drug Effects on Human Induced Pluripotent Stem Cell-Derived Cardiomyocytes,” Toxicological Sciences, vol. 154, no. 2, p. 320, Dec. 2016, doi: 10.1093/TOXSCI/KFW171.

[5] A. Klimas, C. M. Ambrosi, J. Yu, J. C. Williams, H. Bien, and E. Entcheva, “OptoDyCE as an automated system for high-throughput all-optical dynamic cardiac electrophysiology,” Nature Communications, vol. 7, May 2016, doi: 10.1038/NCOMMS11542.

[6] M. Clements, “Multielectrode array (MEA) assay for unit 22.4 profiling electrophysiological drug effects in human stem cell-derived cardiomyocytes,” Current Protocols in Toxicology, vol. 2016, pp. 1–32, May 2016, doi: 10.1002/CPTX.2.

[7] L. Sala, D. Ward-Van Oostwaard, L. G. J. Tertoolen, C. L. Mummery, and M. Bellin, “Electrophysiological Analysis of human Pluripotent Stem Cell-derived Cardiomyocytes (hPSC-CMs) Using Multi-electrode Arrays (MEAs),” Journal of Visualized Experiments : JoVE, vol. 2017, no. 123, p. 55587, May 2017, doi: 10.3791/55587.

[8] H. B. Hayes et al., “Novel method for action potential measurements from intact cardiac monolayers with multiwell microelectrode array technology,” Scientific Reports, vol. 9, no. 1, Dec. 2019, doi: 10.1038/S41598-019-48174-5.

[9] M. Devarasetty et al., “Optical Tracking and Digital Quantification of Beating Behavior in Bioengineered Human Cardiac Organoids,” Biosensors (Basel), vol. 7, no. 3, Jun. 2017, doi: 10.3390/BIOS7030024.

[10] Y. R. Lewis-Israeli et al., “Self-assembling human heart organoids for the modeling of cardiac development and congenital heart disease,” Nature Communications 2021 12:1, vol. 12, no. 1, pp. 1–16, Aug. 2021, doi: 10.1038/s41467-021-25329-5.

[11] J. Suzurikawa, M. Nakao, Y. Jimbo, R. Kanzaki, and H. Takahashi, “A light addressable electrode with a TiO2 nanocrystalline film for localized electrical stimulation of cultured neurons,” Sensors and Actuators, B: Chemical, vol. 192, pp. 393–398, Mar. 2014, doi: 10.1016/j.snb.2013.10.139.

[12] F. Wu et al., “Photoelectrochemical Imaging System for the Mapping of Cell Surface Charges,” 2019, doi: 10.1021/acs.analchem.9b00304.

[13] S. Dantism, S. Takenaga, P. Wagner, T. Wagner, and M. J. Schöning, “Light-addressable potentiometric sensor (LAPS) combined with multi-chamber structures to investigate the metabolic activity of cells,” in Procedia Engineering, Jan. 2015, vol. 120, pp. 384–387. doi: 10.1016/j.proeng.2015.08.647.

[14] J. Wang, L. Du, S. Krause, C. Wu, and P. Wang, “Surface modification and construction of LAPS towards biosensing applications,” Sensors and Actuators, B: Chemical, vol. 265. Elsevier B.V., pp. 161–173, Jul. 15, 2018. doi: 10.1016/j.snb.2018.02.190.

[15] H. Tanaka, T. Yoshinobu, and H. Iwasaki, “Application of the chemical imaging sensor to electrophysiological measurement of a neural cell,” Sensors and Actuators, B: Chemical, vol. 59, no. 1, pp. 21–25, Oct. 1999, doi: 10.1016/S0925-4005(99)00012-X.

[16] A. B. M. Ismail et al., “Investigation on light-addressable potentiometric sensor as a possible cell-semiconductor hybrid,” Biosensors and Bioelectronics, vol. 18, no. 12, pp. 1509–1514, Oct. 2003, doi: 10.1016/S0956-5663(03)00129-5.

[17] Q. Liu et al., “Sensors and Actuators B : Chemical In vitro assessing the risk of drug-induced cardiotoxicity by embryonic stem cell-based biosensor,” Sensors & Actuators: B. Chemical, vol. 155, no. 1, pp. 214–219, 2011, doi: 10.1016/j.snb.2010.11.050.

[18] W. J. Parak et al., “Can the light-addressable potentiometric sensor (LAPS) detect extracellular potentials of cardiac myocytes,” IEEE Transactions on Biomedical Engineering, vol. 47, no. 8, pp. 1106–1113, 2000, doi: 10.1109/10.855939.

[19] J. Wang, Y. Zhou, M. Watkinson, J. Gautrot, and S. Krause, “High-sensitivity light-addressable potentiometric sensors using silicon on sapphire functionalized with self-assembled organic monolayers,” Sensors and Actuators, B: Chemical, vol. 209, pp. 230–236, Mar. 2015, doi: 10.1016/J.SNB.2014.11.071.

[20] J. Wang et al., “The effect of gold nanoparticles on the impedance of microcapsules visualized by scanning photo-induced impedance microscopy,” Electrochimica Acta, vol. 208, pp. 39–46, Aug. 2016, doi: 10.1016/J.ELECTACTA.2016.05.017.

[21] D. W. Zhang, F. Wu, J. Wang, M. Watkinson, and S. Krause, “Image detection of yeast Saccharomyces cerevisiae by light-addressable potentiometric sensors (LAPS),” Electrochemistry Communications, vol. 72, pp. 41–45, Nov. 2016, doi: 10.1016/j.elecom.2016.08.017.

[22] N. Ariyarathna, S. Kumar, S. P. Thomas, W. G. Stevenson, and G. F. Michaud, “Role of Contact Force Sensing in Catheter Ablation of Cardiac Arrhythmias: Evolution or History Repeating Itself?,” JACC: Clinical Electrophysiology, vol. 4, no. 6. Elsevier Inc, pp. 707–723, Jun. 01, 2018. doi: 10.1016/j.jacep.2018.03.014.

[23] A. Thiagalingam et al., “Importance of Catheter Contact Force during Irrigated Radiofrequency Ablation: Evaluation in a Porcine *Ex Vivo* Model Using a Force-Sensing Catheter,” Journal of Cardiovascular Electrophysiology, vol. 21, no. 7, pp. 806–811, Jan. 2010, doi: 10.1111/j.1540-8167.2009.01693.x.

[24] “TactiCath Quartz Contact Force Ablation Catheter | Abbott.” https://www.cardiovascular.abbott/us/en/hcp/products/electrophysiology/tacticath-quartz-contact-force-ablation-catheter.html (accessed Apr. 23, 2020).

[25] R. Cornell, “Cell-substrate adhesion during cell culture: An ultrastructural study,” Experimental Cell Research, vol. 58, no. 2-3, pp. 289–295, Dec. 1969, doi: 10.1016/0014-4827(69)90507-2.

[26] E. Kreysing, H. Hassani, N. Hampe, and A. Offenhäusser, “Nanometer-Resolved Mapping of Cell-Substrate Distances of Contracting Cardiomyocytes Using Surface Plasmon Resonance Microscopy,” ACS Nano, vol. 12, no. 9, pp. 8934–8942, Sep. 2018, doi: 10.1021/ACSNANO.8B01396.

[27] D. Zhao, W. Lei, and S. Hu, “Cardiac organoid — a promising perspective of preclinical model,” Stem Cell Research and Therapy, vol. 12, no. 1, pp. 1–10, Dec. 2021, doi: 10.1186/S13287-021-02340-7/FIGURES/3.

[28] L. Chen, Y. Zhou, S. Jiang, J. Kunze, P. Schmuki, and S. Krause, “High resolution LAPS and SPIM,” Electrochemistry Communications, vol. 12, no. 6, pp. 758–760, Jun. 2010, doi: 10.1016/j.elecom.2010.03.026.

[29] S. Scalzo et al., “Dense optical flow software to quantify cellular contractility,” Cell Reports Methods, vol. 1, no. 4, p. 100044, Aug. 2021, doi: 10.1016/J.CRMETH.2021.100044.

[30] A. F. Kalmar et al., “Phenylephrine increases cardiac output by raising cardiac preload in patients with anesthesia induced hypotension,” J Clin Monit Comput, vol. 32, no. 6, pp. 969–976, Dec. 2018, doi: 10.1007/S10877-018-0126-3.

[31] E. Richards, M. J. Lopez, and C. v. Maani, “Phenylephrine,” xPharm: The Comprehensive Pharmacology Reference, pp. 1–5, Jul. 2021, doi: 10.1016/B978-008055232-3.62411-0.

[32] M. Gálvez-Santisteban et al., “Hemodynamic-mediated endocardial signaling controls in vivo myocardial reprogramming,” Elife, vol. 8, Jun. 2019, doi: 10.7554/ELIFE.44816.

[33] A. Brodarac et al., “Susceptibility of murine induced pluripotent stem cell-derived cardiomyocytes to hypoxia and nutrient deprivation,” Stem Cell Res Ther, vol. 6, no. 1, Apr. 2015, doi: 10.1186/S13287-015-0057-6.

[34] J. Y. Min et al., “Transplantation of embryonic stem cells improves cardiac function in postinfarcted rats,” Journal of Applied Physiology, vol. 92, no. 1, pp. 288–296, 2002, doi: 10.1152/JAPPL.2002.92.1.288/ASSET/IMAGES/LARGE/DG0121216107.JPEG.

[35] A. O. Verkerk, C. C. Veerman, J. G. Zegers, I. Mengarelli, C. R. Bezzina, and R. Wilders, “Patch-Clamp Recording from Human Induced Pluripotent Stem Cell-Derived Cardiomyocytes: Improving Action Potential Characteristics through Dynamic Clamp,” International Journal of Molecular Sciences, vol. 18, no. 9, Sep. 2017, doi: 10.3390/IJMS18091873.

[36] L. M. Hondeghem, L. Carlsson, and G. Duker, “Instability and triangulation of the action potential predict serious proarrhythmia, but action potential duration prolongation is antiarrhythmic,” Circulation, vol. 103, no. 15, pp. 2004–2013, Apr. 2001, doi: 10.1161/01.CIR.103.15.2004.

[37] F. Li et al., “Inhibition of myosin IIA–actin interaction prevents ischemia/reperfusion induced cardiomyocytes apoptosis through modulating PINK1/Parkin pathway and mitochondrial fission,” International Journal of Cardiology, vol. 271, pp. 211–218, Nov. 2018, doi: 10.1016/J.IJCARD.2018.04.079.

[38] P. Pandey et al., “Cardiomyocytes Sense Matrix Rigidity through a Combination of Muscle and Non-muscle Myosin Contractions,” Dev Cell, vol. 45, no. 5, p. 661, Jun. 2018, doi: 10.1016/J.DEVCEL.2018.05.016.

[39] B. Zhou et al., “Photoelectrochemical imaging system with high spatiotemporal resolution for visualizing dynamic cellular responses,” Biosensors and Bioelectronics, vol. 180, p. 113121, May 2021, doi: 10.1016/J.BIOS.2021.113121.

[40] R. W. Schafer, “What is a savitzky-golay filter?,” IEEE Signal Processing Magazine, vol. 28, no. 4, pp. 111–117, 2011, doi: 10.1109/MSP.2011.941097.

