## Supporting Information for "Photoelectrochemical imaging of action potentials for single cardiomyocytes through contact force manipulation of organoids"

**Line scans of organoids**

For the organoids several line scans (Figure S1b) are completed at the same location and from this 'image' of line scans (Figure S1a) a time dependent photocurrent at each X position along the line can be established (Figure S1c). Each line takes 0.1 s to complete and so the photocurrent is sampled at each location with 0.1 s intervals. From the line scan image, it is clear that measurements at the edges of the cells are subject to changes due to the mechanical movement of the cell; for this reason, all photocurrent vs time cross sections are taken at the centre of each cell. All line scans contain 80 pixels but vary in length according to the size, number and orientation of the cells measured in each case.

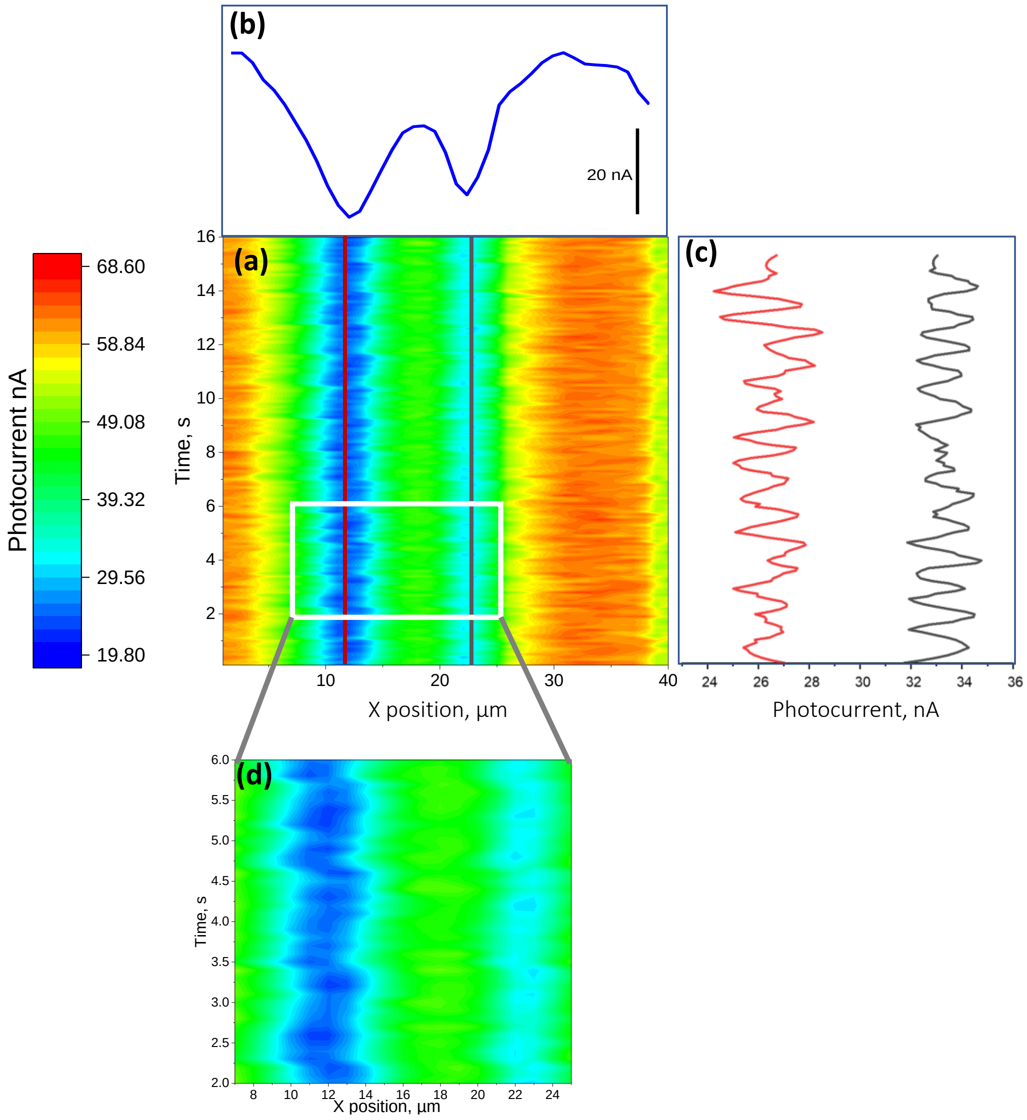


*Figure S1*
*Line scans: (a) A line is continually scanned to build up photocurrent image (b). Horizontal lines across the image indicate the position of cells by minima in the photocurrent (c) vertical cross sections are taken at the centre of each cell present producing corresponding photocurrent vs time data. (d) zoomed in view of line scan image showing movement at the edge of each cell*

### Smoothing of photocurrent vs time curves

The raw measured photocurrent data (figure S2a) has a large amount of high frequency noise; hence, a smoothing method was implemented to filter out the noise. The method used was the Savitzky-Golay method which is able to preserve peak profiles within the data [40] (Figure S2b)

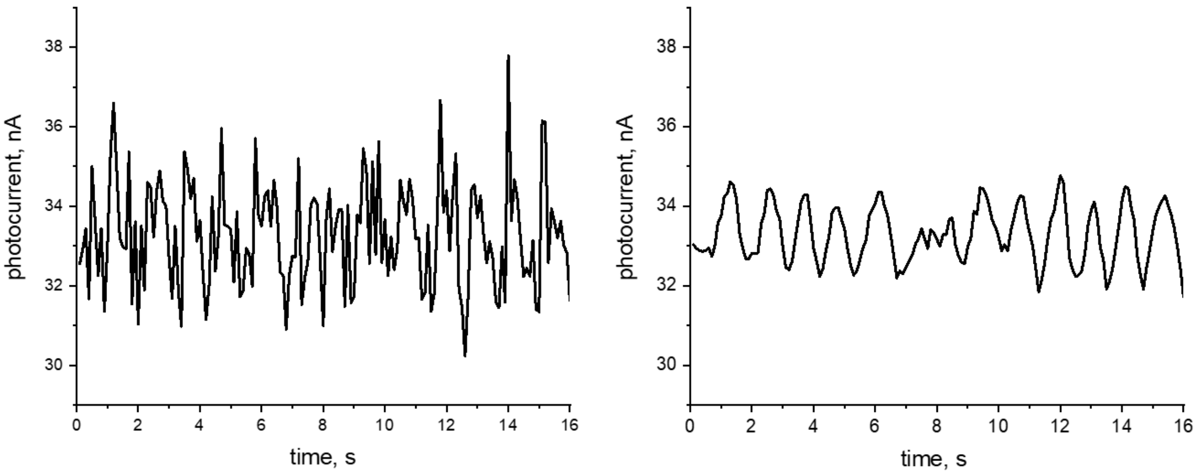


*Figure S2*
(a*) The raw photocurrent vs time signal for a cell (b) corresponding smoothed photocurrent from using the Savitzky-Golay method (10 pts)*

### Phase determination through sinusoidal fit

The phase shift of each photocurrent time dependence is determined through the fitting of a sine curve. The curve of best fit is determined in origin pro with the value of phase shift noted for each. A phase difference can then be obtained for the two adjoining cells. Figure S3 describes the sine wave fitted to each cell's photocurrent response for before and after verapamil is added. The sinusoidal formula is noted and the parameters of each sine wave is detailed in the Table S1; the xc values are highlighted, and errors from the fitting are also noted.

A phase difference for cell 1 and cell 2 before and after verapamil is added can now be determined by simply converting the phase shift in seconds to degrees using the value of beat period (formula S1) and finding the difference. In this example there is a phase difference of 165 ± 11 ° before the verapamil is added and 7.37 ± 11 ° afterward. The error in phase difference is determined by the standard error propagation. The phase difference drug induced changes are determined for all adjoining cells and are detailed in Tables S3 and S4.

$$\boldsymbol{P}^{\circ} = \left(\frac{\boldsymbol{P_s}}{\boldsymbol{T}}\right) \boldsymbol{x\ 360}$$

*Equation S1 conversion formula where P is phase in degrees P is phase in s and T is the beat period*

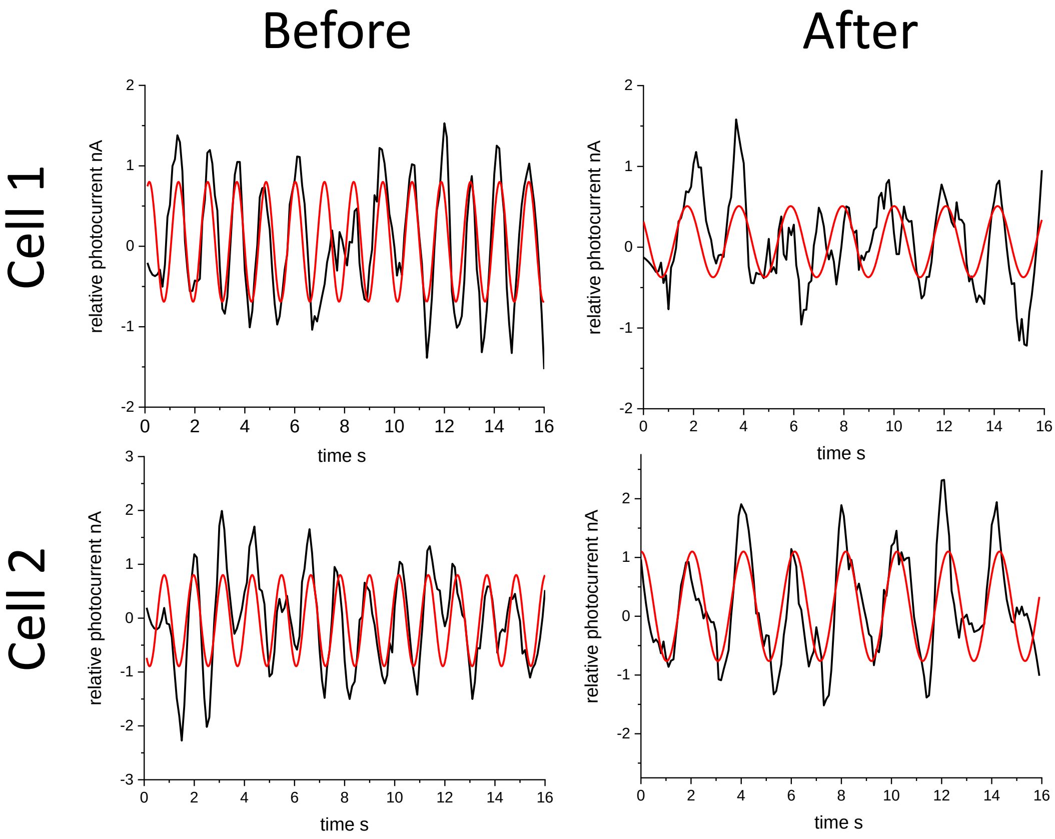


$$y = y0 + A\sin\left(\pi\frac{(x-xc)}{w}\right)$$

*Figure S3*
*the phase determination method, sine wave fitting to the photocurrent time dependence measured for cell 1 and cell 2 before and after verapamil is added and the sinusoidal formula where y0 is the y intercept, xc is the phase shift, A the amplitude and w is the period*

*Table S1 the parameters of the sine wave fitting detailed in Figure S4, highlighting the phase shift xc values with errors in the fit*

| **Parameter** | **Cell 1** | | **Cell 2** | |
|---|---|---|---|---|
| | before | after | before | after |
| y0 nA | -0.0453 | 0.0551 | 0.0557 | 0.170 |
| xc s | 0.507 ± 0.028 | -0.432 ±0.044 | -0.0898 ±0.27 | -0.389 ± 0.045 |
| w s | 0.585 | 1.03 | 0.585 | 1.02 |
| A nA | 0.847 | 0.440 | 0.745 | 0.931 |

**Duration parameters**

The duration parameters are determined from several APs as defined in figure S4. The average value for each parameter before and after each drug is detailed in Table S2 along with the corresponding statistical significance.

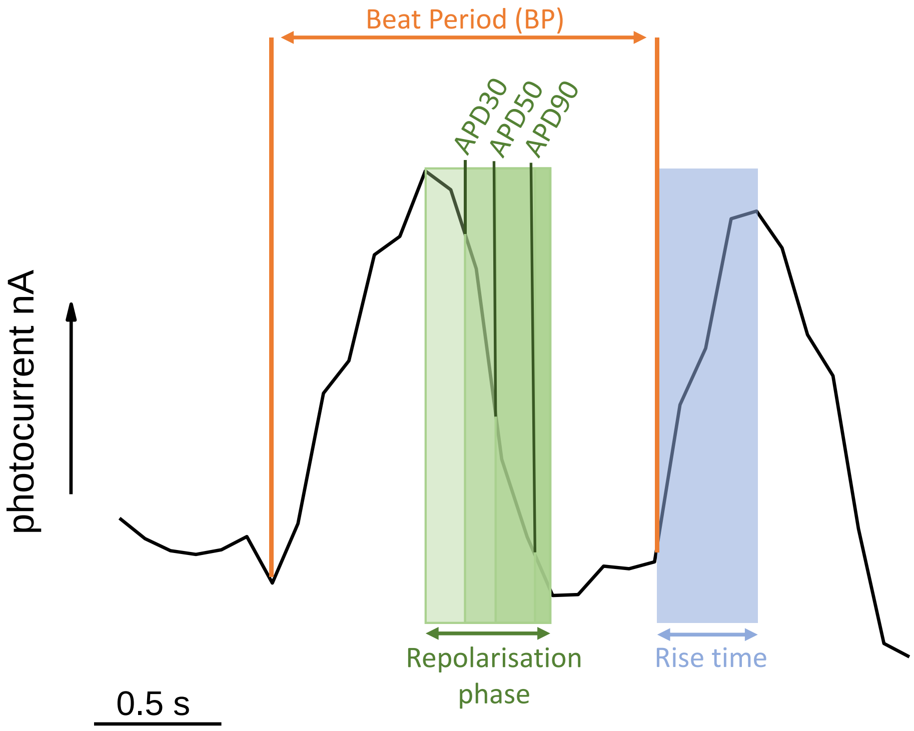


*Figure S4*
*Duration parameter definitions, shows how the beat period, rise time and APD30, 50 and 90 are determined for each AP using an example photocurrent vs time measurement.*

The parameters determined for both control and drugs are listed in Table S2 with stars corresponding to the p-value determined for the change in mean, where *(p<0.05) **(p<0.01) and ***(p<0.001)

*Table S2 the average duration parameters before and after each drug statistically significant changes are highlighted with *s according to their p value*

| | | Before | SEM | After | SEM |
|---|---|---|---|---|---|
| Verapamil | Rise Time***, s | 0.504 | 0.0178 | 0.692 | 0.036 |
| | BP***, s | 1.04 | 0.03 | 1.52 | 0.06 |
| | APD30***, s | 0.706 | 0.0195 | 0.926 | 0.038 |
| | APD50***, s | 0.794 | 0.016 | 1.16 | 0.05 |
| | APD90***, s | 0.981 | 0.027 | 1.42 | 0.06 |
| | Triangulation | 0.818 | 0.018 | 0.820 | 0.020 |
| | Amplitude*, nA | 1.00 | 0.03 | 0.891 | 0.045 |
| PE | Rise Time**, s | 0.810 | 0.064 | 0.610 | 0.032 |
| | BP*,s | 1.48 | 0.096 | 1.13 | 0.03 |
| | APD30, s | 1.07 | 0.09 | 0.965 | 0.075 |
| | APD50, s | 1.15 | 0.09 | 1.04 | 0.08 |
| | APD90, s | 1.42 | 0.10 | 1.21 | 0.08 |
| | Triangulation | 0.810 | 0.022 | 0.849 | 0.016 |
| | Amplitude*, nA | 1.00 | 0.05 | 1.17 | 0.06 |
| Blebbistatin | Rise Time, s | 0.415 | 0.031 | 0.255 | 0.017 |
| | BP, s | 1.02 | 0.04 | 0.935 | 0.040 |
| | APD30, s | 0.623 | 0.027 | 0.577 | 0.029 |
| | APD50, s | 0.716 | 0.027 | 0.671 | 0.029 |
| | APD90, s | 0.959 | 0.041 | 0.858 | 0.033 |
| | Triangulation | 0.751 | 0.018 | 0.782 | 0.017 |
| | Amplitude, nA | 1.00 | 0.10 | 0.995 | 0.102 |

**Data tables**

Table S3 details the data for cardiomyocytes cultured on the LAPS sensor surface noting the average amplitude of each cell and approximate frequencies, determined through fitting a sine wave to the photocurrent vs time data.

The change in the phase of the APs acquired before and after adding verapamil, PE and blebbistatin are detailed in table S4, S5 and S6 respectively, including, in the case where two cells are adjoining, the phase shift. The raw amplitude averages are also recorded with SEM. In the case of verapamil, the phase difference is reduced and in the case of PE little difference to the phase is observed.

*Table S3 the data analysis acquired from monitoring the photocurrent individual cells cultured on the sensor surface*

| culture | Cell measured | Amplitude nA | SE amplitude ± nA | Frequency, Hz | Error in frequency ± Hz |
|---|---|---|---|---|---|
| 1 | 1 | 0.595 | 0.0594 | 1.27 | 0.0143 |
| | 2 | 0.318 | 0.0459 | 2.09 | 0.0631 |
| | 3 | 0.427 | 0.0783 | 2.72 | 0.0126 |
| 2 | 4 | 0.454 | 0.133 | 1.64 | 0.0277 |
| | 5 | 0.227 | 0.0787 | 1.26 | 0.0280 |
| | 6 | 0.141 | 0.0210 | 1.66 | 0.0213 |

*Table S4 the data analysis acquired from monitoring the photocurrent of cardiomyocyte cells before and 10 mins after the addition of verapamil*

| Organoid number | Cell number | | Amplitude at start nA | Phase shift s | Phase difference between adjoining cells, ° | |
|---|---|---|---|---|---|---|
| | | | | | before | after |
| 1 | 1 | Before | 2.25 (±0.231) | 0.507 (± 0.028) | 165 (± 11) | 7.37 (± 10.7) |
| | | After | 1.35 (±0.170) | -0.432 (± 0.044) | | |
| | 2 | Before | 2.68 (±0.246) | -0.0896 (± 0.0273) | | |
| | | After | 1.99 (±0.128) | -0.389 (± 0.045) | | |
| 2 | 3 | Before | 0.975 (±0.126) | 0.784 (± 0.103) | 64.7 (± 38.3) | 2.22 (± 22.56) |
| | | After | 0.900 (±0.132) | -0.350 (± 0.022) | | |
| | 4 | Before | 0.842 (±0.0728) | 0.541 (±0.100) | | |
| | | After | 0.810 (±0.0869) | --0.362 (±0.120) | | |
| 3 | 5 | Before | 0.854 (±0.0924) | | | |
| | | After | 0.771 (±0.0820) | | | |

*Table S5 the data analysis acquired from monitoring the photocurrent of cardiomyocyte cells before and 2-5mins after the addition of PE*

| Organoid number | Cell number | | Average amplitude, nA | Phase shift | Phase difference between adjoining cells, ° | |
|---|---|---|---|---|---|---|
| | | | | | before | after |
| 4 | 6 | Before | 0.579 (±0.054) | | | |
| | | After | 0.610 (±0.066) | | | |
| 5 | 7 | Before | 1.08 (±0.14) | 1.11 (± 0.10) | 66 ( ± 55) | 56 (± 52) |
| | | After | 1.34 (±0.12) | 0.738 (± 0.096) | | |
| | 8 | Before | 1.45 (±0.17) | 1.25 (± 0.06) | | |
| | | After | 1.72 (±0.18) | 1.33 (± 0.07) | | |
| 6 | 9 | Before | 1.01 (±0.16) | | | |
| | | After | 1.17 (±0.15) | | | |

*Table S6 the data analysis acquired from monitoring the photocurrent of cardiomyocyte cells before and 2-5mins after the addition of blebbistatin*

| Organoid number | Cell number | | Average amplitude, nA |
|---|---|---|---|
| 7 | 10 | Before | 1.70 (±0.08) |
| | | After | 1.56 (±0.20) |
| | 11 | Before | 1.74 (±0.28) |
| | | After | 1.41 (±0.10) |
| 8 | 12 | Before | 0.572 (±0.058) |
| | | After | 0.535 (±0.071) |
| | 13 | Before | 0.492 (±0.025) |
| | | After | 0.476 (±0.066) |
